# Thermal characterization and interaction of the subunits from the multimeric bacteriophage endolysin PlyC

**DOI:** 10.1101/2022.01.06.475266

**Authors:** J. Todd Hoopes, Ryan D. Heselpoth, Frederick P. Schwarz, Daniel C. Nelson

## Abstract

Bacteriophage endolysins degrade the bacterial peptidoglycan and are considered enzymatic alternatives to small molecule antibiotics. In particular, the multimeric streptococcal endolysin PlyC has appealing antibacterial properties. However, a comprehensive thermal analysis of PlyC is lacking, which is necessary for evaluating long-term stability and downstream therapeutic potential. Biochemical and kinetic-based methods were used in combination with differential scanning calorimetry to investigate the structural, kinetic and thermodynamic stability of PlyC and its various subunits and domains. The PlyC holoenzyme structure is irreversibly compromised due to partial unfolding and aggregation at 46°C. Unfolding of the catalytic subunit, PlyCA, instigates this event, resulting in the kinetic inactivation of the endolysin. In contrast to PlyCA, the PlyCB octamer (the cell wall binding domain) is thermostable, denaturing at ∼75°C. Isolation of PlyCA or PlyCB alone altered their thermal properties. Contrary to the holoenzyme, PlyCA alone unfolds uncooperatively and is thermodynamically destabilized whereas the PlyCB octamer reversibly dissociates into monomers and forms an intermediate state at 74°C in phosphate buffered saline, with each subunit subsequently denaturing at 92°C. Adding folded PlyCA to an intermediate state PlyCB, followed by cooling, allowed for *in vitro* reconstitution of the active holoenzyme.

## Introduction

The increasing global threat of bacterial resistance to the traditional antibiotics highlights the need for alternative antimicrobials (1, 2). One potential antimicrobial strategy involves the use of bacteriophage (or phage)-encoded endolysins (3, 4). These enzymes are primarily peptidoglycan (i.e., cell wall) hydrolases that cleave critical covalent bonds in the cell wall of phage-infected bacteria. Due to the elevated intracellular turgor pressure, the loss of peptidoglycan structural integrity causes hypotonic lysis and phage virion liberation. The antibacterial efficacy of extrinsically-applied recombinant endolysins toward sensitive bacteria has been validated both *in vitro* and *in vivo* (5, 6).

One example of an endolysin with established antibacterial activity of clinical interest is PlyC. This endolysin rapidly kills species of *Streptococcus* that cause both human and animal disease (7, 8). Interestingly, PlyC eradicates *Streptococcus pyogenes* biofilms and innately possesses the unique ability to translocate across epithelial cell membranes to kill intracellular streptococci (9, 10). In addition to its potent *in vitro* antibacterial properties, PlyC was highly effective *in vivo* when administered as either a prophylactic or therapeutic using a murine pharyngeal decolonization model (7).

The structure of PlyC has a novel multimeric composition (11). Contrary to the conventional protein architecture for endolysins that target Gram-positive bacteria, which generally consist of a single polypeptide with a conserved N-terminal catalytic domain linked to a variable C-terminal cell wall binding domain (CBD), the PlyC holoenzyme collectively consists of nine subunits. The CBD comprises eight identical PlyCB monomers that interact through protein-protein interactions to form an octameric ring structure (11). The PlyCB CBD was recently shown to bind with high affinity and avidity to a polyrhamnose moiety shared in the wall teichoic acids of sensitive streptococcal species (12, 13). The ninth subunit, PlyCA, is the catalytic subunit consisting of an N-terminal glycosyl hydrolase (GyH) domain, a central helical docking domain, and a C-terminal cysteine, histidine-dependent amidohydrolase/peptidase (CHAP) domain. The GyH and CHAP domains act together synergistically to generate the robust bacteriolytic activity of PlyC, whereas the central helical docking domain of PlyCA interacts with the PlyCB CBD to form the holoenzyme structure (11).

Elucidating the thermal characteristics of a biologic system is necessary for assessing its long-term stability and therapeutic potential. Moreover, the Arrhenius equation describes the relationship between the rate of a chemical reaction and temperature, and thereby associates the thermal stability of a material in general, and a drug in particular, to its shelf-life expectancy. Accordingly, at a given temperature, the intrinsic stability of a protein directly correlates with the activation energy required for unfolding and, by extension, shelf-life (14, 15). With this in mind, we determined the thermal properties of PlyC. Due to the complexity of its multimeric structure, the PlyC holoenzyme, as well as individual subunits and domains, were thermally characterized using a combination of biological, functional and biophysical experimental approaches. Results from this analysis will ultimately aid in downstream developmental strategies for prolonging long-term stability of PlyC, such as identifying thermolabile structural components to target in subsequent protein engineering studies and excipient formulations.

## Materials and methods

### Bacterial strains, culture conditions, expression and purification

1. *S. pyogenes* strain D471 was used for all experiments and routinely grown in Todd Hewitt broth supplemented with 1% (w/v) yeast extract as previously described (7, 16). *Escherichia coli* strains used for protein expression were grown at 37°C in 4 L baffled flasks containing 1.5 L of Luria-Bertani and ampicillin (100 µg/ml when needed) with shaking at 190 revolutions per minute. Recombinant PlyC, PlyCA, PlyCB and CHAP were expressed and purified as previously described (11).

### 15-day kinetic stability assay

PlyC at 1 mg/ml was incubated at either 4, 25 or 37°C in phosphate buffered saline (PBS), pH 7.2, for a total of 15 days. In 48 h increments, an aliquot from three independent PlyC samples at each temperature was assayed in triplicate for residual lytic activity at a final concentration of 5 μg/ml towards *S. pyogenes* strain D471 using a turbidity reduction assay. For this, an overnight culture of *S. pyogenes* were harvested, washed and resuspended in PBS to an optical density at 600 nm (OD_600nm_) of 2.0. Equal volumes (100 µl each) of the temperature-treated PlyC samples and *S. pyogenes* were mixed in individual wells of a 96-well flat-bottomed microtiter plate. Using a SpectraMax 190 spectrophotometer (Molecular Devices), the OD_600nm_ was measured every 15 sec for a total of 20 min at 37°C. Activity was equated to the V_max_ (milli-OD units per min) of the resulting curve corresponding to the loss in optical density over time. Data was averaged and normalized to the bacteriolytic activity displayed by each PlyC sample on day 1.

### Thermal denaturation of PlyC

PlyC aliquots at 1 mg/ml were incubated in phosphate buffered saline (PBS), pH 7.2, using EchoTherm™ heat blocks (Torrey Pines Scientific) at 37, 42, 50, 55, 60 or 65°C for 30 min. The samples were then removed and immediately cooled on ice. The structural integrity of the holoenzyme was assessed using native polyacrylamide gel electrophoresis (PAGE). Sodium dodecyl sulfate polyacrylamide gel electrophoresis (SDS-PAGE) was used as a control to confirm the presence of both the PlyCA and PlyCB subunits at each temperature. Once the temperature range for thermal denaturation was narrowed, the experiment was repeated, but at temperatures of 42, 43, 44, 45, 46, 47 or 48°C.

The kinetic stability of PlyC was additionally assayed at each temperature described above. For these experiments, PlyC at 1 mg/ml in PBS was heated for 10 min using an EchoTherm^TM^ heat block and then immediately cooled on ice. Turbidity reduction assays were then used to measure residual bacteriolytic activity. For these experiments, an overnight culture of *S. pyogenes* strain D471 were harvested, washed and resuspended in PBS to an OD_600nm_ of 1.0. Equal volumes (100 µl each) of the heated PlyC samples and bacteria were mixed into individual wells of a 96-well flat-bottomed microtiter plate. Using a SpectraMax 190 spectrophotometer, the OD_600nm_ was measured every 10 sec for a total of 5 min at room temperature. Bacteriolytic activity was equated to the difference in OD_600nm_ over the duration of the experiment. All data was normalized to the activity of PlyC without heat treatment.

### Differential scanning calorimetry

Differential scanning calorimetry (DSC) measurements were performed using a VP-DSC Microcalorimeter (MicroCal Inc.) under a static, oxidative atmosphere. The sample and reference vessels had an operational volume of 0.511 ml. Prior to sample analysis, PBS, pH 7.4, or 20 mM phosphate buffer, pH 7.4, was initially added to both the sample and reference vessels and subsequently subjected to two heating/cooling cycles from 15 to 105°C and 105 to 15°C using scan rates of either 15 or 60 K/h. The result from the second heating/cooling cycle was utilized as the base line. Next, protein samples at various concentrations were independently injected into the sample vessel and subjected to multiple heating/cooling cycles. After subtraction of the baseline from the sample scan, the resulting raw data of the differential power as a function of time was divided by the scan rate to convert the data into heat capacity versus temperature. Utilizing the EXAM program (17), two-state thermal transition models were fitted to the heat capacity versus temperature data to calculate the van’t Hoff enthalpy (Δ*H*_VH_) from the shape of the transition peak. EXAM was also used to determine *T*_G_ (the temperature at the thermal transition midpoint) and calorimetric enthalpy (Δ*H*_cal_; the area under the transition peak divided by the moles of protein in the sample cell) values.

### PlyCB cross-linking

At 1 mg/ml, 100 μl aliquots of the PlyCB CBD in PBS were heated for 10 min in a gradient thermocycler (MyCycler, BioRad) at temperatures ranging from 25 to 90°C. Following the instructions provided by the manufacturer, the amine groups of the heated PlyCB samples were crosslinked with bis(sulfosuccinimidyl) suberate (ThermoFisher Scientific) for 10 min. After the reaction was quenched with 10 µl of 1 M Tris, pH 7.4, the heated samples were immediately placed on ice for 10 min and ultimately resolved via SDS-PAGE.

### Bright field and fluorescence microscopy

The organic dye, AlexaFluor™-555 (ThermoFisher Scientific), was conjugated to PlyCB according to the manufacturer’s instructions. Aliquots consisting of 100 µl of the conjugated protein at 1 mg/ml were incubated from 4 to 100°C for 30 min in PBS using an EchoTherm™ refrigerated heat block and then immediately cooled on ice. Next, the protein samples (10 µl each) were incubated with *S. pyogenes* in PBS for 10 min at room temperature. The labeled cells were washed with PBS and viewed on a Nikon Eclipse 80i fluorescent microscope equipped with a Retiga 2000R camera using a Cy3 filter for fluorescence imaging.

### Thermal regeneration of native PlyC from its constituent subunits

At a concentration of 15.6 μM (1 mg/ml), 500 μl of the PlyCB CBD in PBS was placed in an EchoTherm™ heat block and heated to 75°C. Next, 500 μl of 19.5 μM (1 mg/ml) ice-cold PlyCA was rapidly added via a syringe to the heated PlyCB sample. Alternately, an identical experiment was carried out with both PlyCA and PlyCB protein samples at room temperature. Immediately after mixing, all samples were placed in an ice bath and allowed to cool for 5 min. Samples were tested for residual lytic activity against *S. pyogenes* using a turbidity reduction assay as previously described (see Kinetic inactivation assay). Relative lytic activity was normalized to that of wild-type PlyC, and equal molar concentrations of isolated PlyCA or PlyCB CBD were used as controls. To further establish that the PlyC holoenzyme had been reconstituted by mixing heated PlyCB with cold PlyCA, the remaining sample was centrifuged at 16,000 x *g* for 10 min to remove any precipitate and loaded onto a HiPrep 16/60 Sephacryl S-200 HR size-exclusion column (GE Healthcare) pre-equilibrated in PBS. An elution peak corresponding to native PlyC was collected and subsequently analyzed by SDS-PAGE.

## Results

### Kinetic stability and thermal denaturation of PlyC

The long-term kinetic stability of PlyC was assessed over an expanded temperature range consisting of 4, 25, and 37°C. The residual bacteriolytic activity of the endolysin was measured against *S. pyogenes* in 48 h increments over a total of 15 days. Findings from these studies indicate PlyC was stable at 4 and 25°C, with the endolysin respectively retaining 100% and 81% residual activity over the duration of the experiment (Fig. 1). Conversely, PlyC was unstable at 37°C. The endolysin retained only 33% activity by day nine and was largely inactivated at the culmination of the experiment.

**Figure 1.**
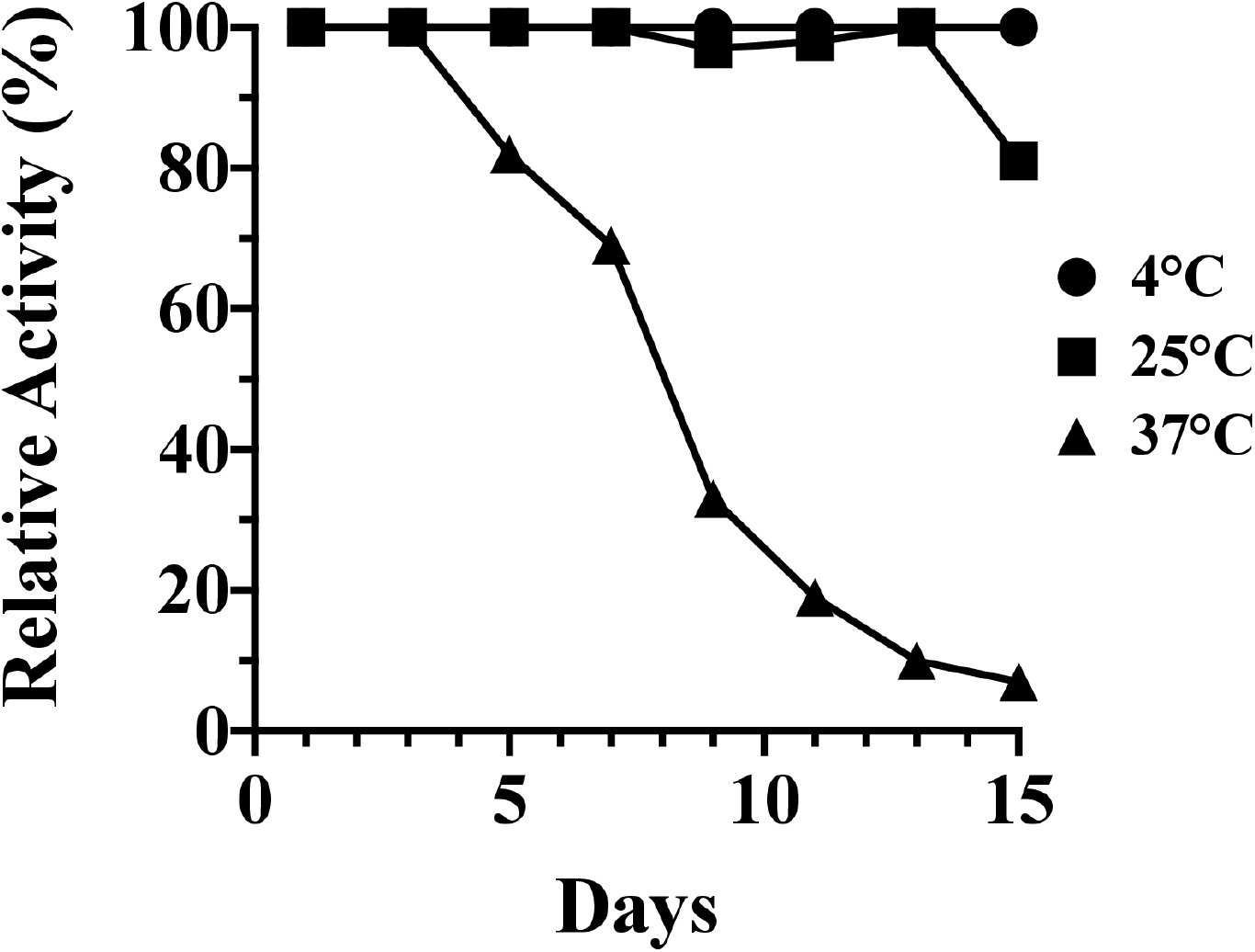
15-day kinetic stability analysis of PlyC. The kinetic stability of PlyC at 1 mg/ml in PBS, pH 7.2, was evaluated for 15 days at 4, 25 and 37°C. At each temperature, an aliquot from three independent samples of PlyC was removed every 48 h and subsequently assayed for residual bacteriolytic activity towards *S. pyogenes* strain D471 using a turbidity reduction assay. Data was averaged and normalized to the bacteriolytic activity of each sample observed on day 1.

Short-term heating experiments were utilized next to identify the precise temperature PlyC is thermally denatured. For these studies, the thermal stability of the PlyC holoenzyme structure was first examined by incubating equal molar concentrations of the purified endolysin at temperatures ranging from 37 to 65°C for 30 minutes, followed by rapid cooling and visual inspection using native PAGE (Fig. 2). Results from the initial electrophoretic analysis suggest the structural conformation of PlyC is compromised between 42 and 50°C, as evidenced by the inability of the aggregated sample to enter a native polyacrylamide gel (Fig. 2, left). When samples at all temperatures were analyzed on an SDS-PAGE gel, the PlyCA and PlyCB subunits were observed at relatively equal concentrations (data not shown), thus proving the absence of bands on native PAGE for PlyC treated at ≥ 50°C can be directly attributed to protein unfolding and aggregation. Next, PlyC was heated from 42 to 48°C in one degree increments in order to determine the precise temperature the holoenzyme loses structural integrity. Native PAGE revealed PlyC loses electrophoretic mobility at 46°C, as a result of thermally-induced aggregation (Fig. 2, right). These observations directly correlate with a temperature-dependent loss of enzymatic activity toward *S. pyogenes* (Fig. 2, bottom).

**Figure 2.**
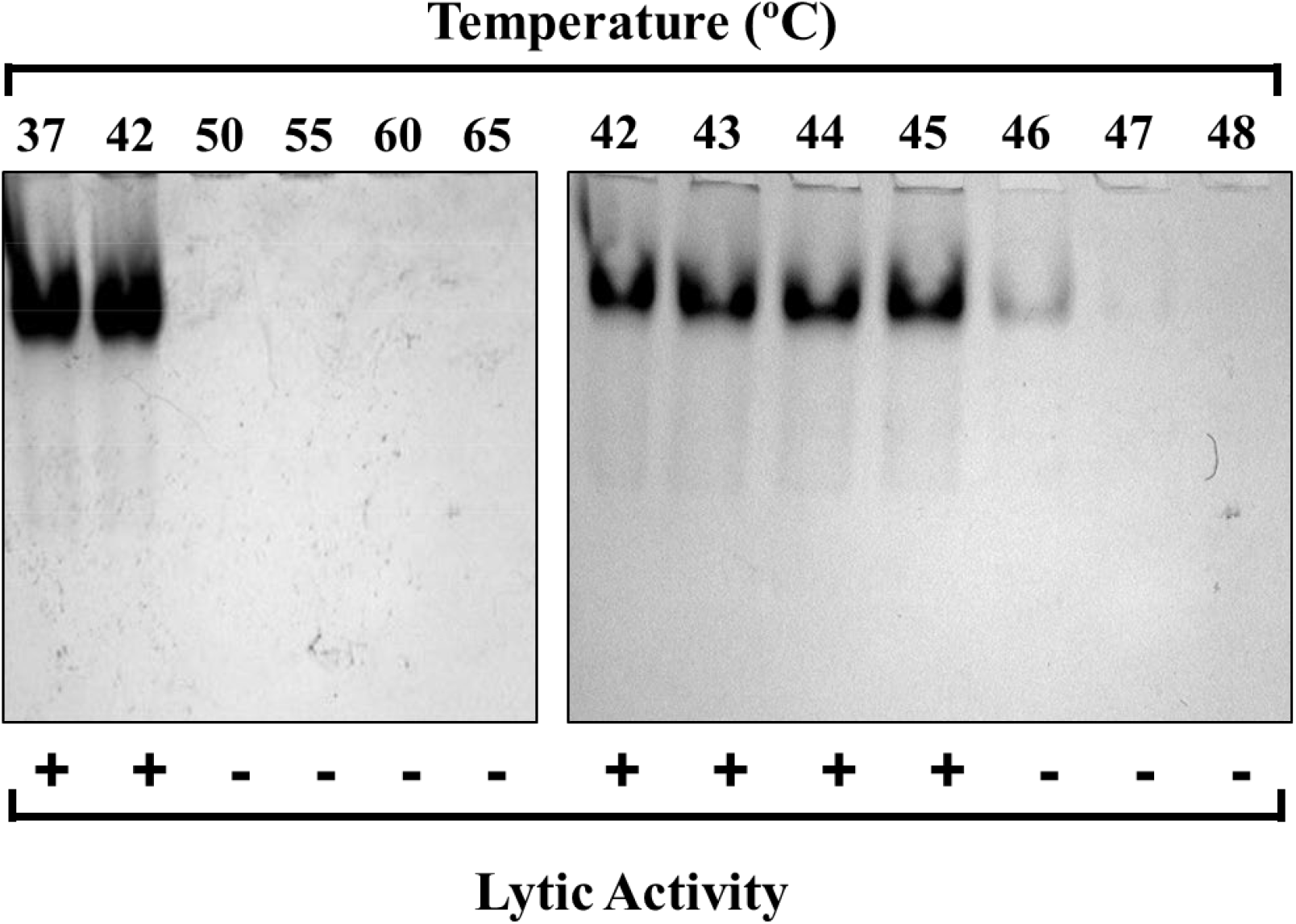
Thermal denaturation of PlyC. Top: The structural stability of the PlyC holoenzyme was assessed by incubating the endolysin at 1 mg/ml in PBS at temperatures ranging from 37 to 65°C (left) o 42 to 48°C (right) for 30 min. After heat treatment, the samples were cooled on ice and visualized via native PAGE. Absence of PlyC in wells corresponds to thermally-induced protein unfolding and aggregation. Bottom: Lytic activity was normalized to unheated PlyC. +, ≥ 90% of unheated PlyC activity; -, ≤ 10% of unheated PlyC activity.

### Thermodynamic characterization of PlyC

DSC was performed to investigate the thermodynamic properties of the PlyC holoenzyme, as well as various isolated subunits and domains of the enzyme. Each protein sample investigated was purified to near homogeneity (Fig. 3). For this thermodynamic analysis, thermal transitions detected for PlyCA-related structural components were fit with A ⟷ I and I ⟷ B two-state transition models specific to the heat capacity versus temperature data. Thermal transitions for the PlyCB octamer were fit with either 2A ⟷ 2I and 2I ⟷ 2B two-state transition models (for the isolated CBD) or a 2A ⟷ 2B two-state transition model (in the context of the PlyC holoenzyme). The resulting *T_G_* and enthalpy values obtained for each symmetrical or asymmetrical thermal denaturation transition corresponds to the two-state transition model that fit best. Transition properties that are considered to be scan rate independent, which can be concluded when the *T_G_* values with twice their uncertainties are within 3°C and the enthalpies are within twice their uncertainties, are capable of being analyzed by thermodynamic two-state transition models (18).

**Figure 3.**
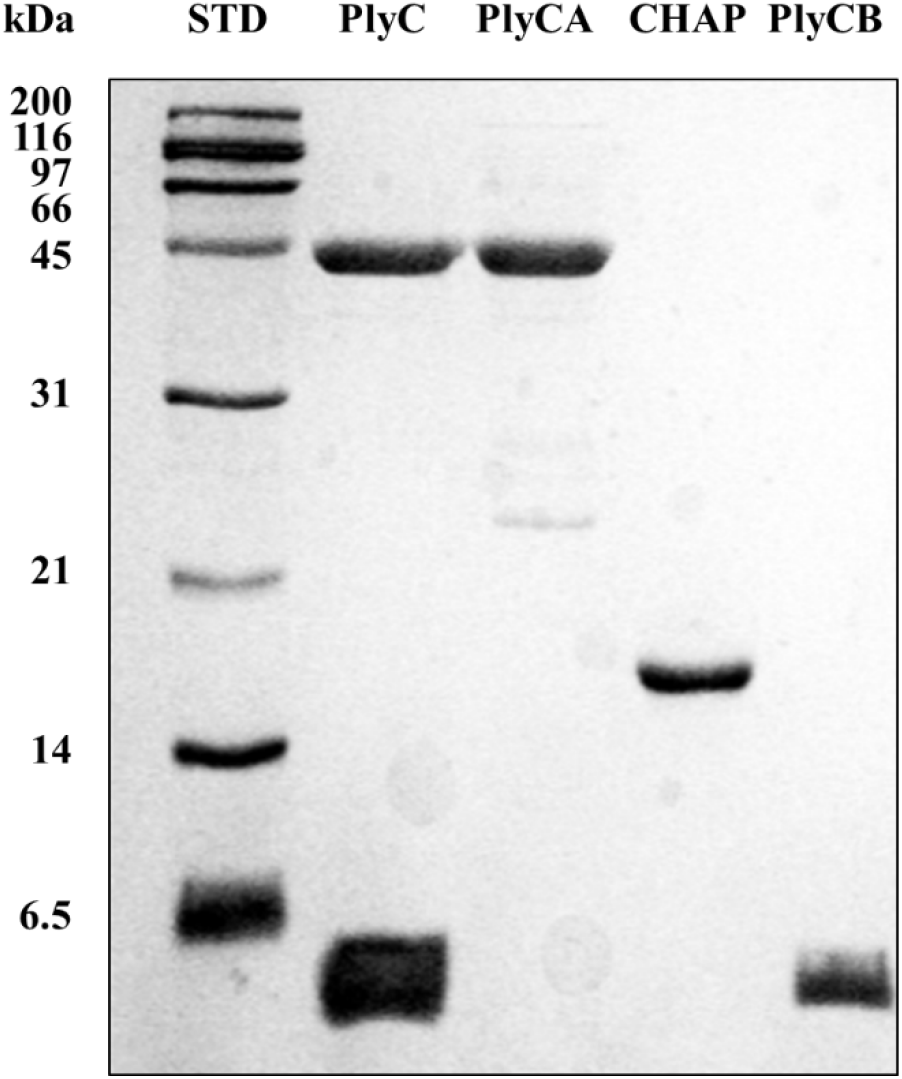
**Purification of various PlyC-related constructs used in this study**. The PlyC holoenzyme (114 kDa; PlyCA subunit = 50 kDa, PlyCB subunit = 8 kDa), the PlyCA catalytic subunit (50 kDa), the C-terminal CHAP domain of PlyCA (18 kDa), and the PlyCB CBD (64 kDa; PlyCB subunit = 8 kDa) were purified to near homogeneity as indicated by SDS-PAGE.

First, PlyCA was thermodynamically characterized at 1 to 2 mg/ml in PBS using scan rates of 15 and 60 K/h (Table 1). As an example, the DSC thermogram of PlyCA is shown in Fig. 4A. DSC scans of PlyCA were fit with A ⟷ I and I ⟷ B two-state transitions and thus fulfills a three-state thermal transition model summarized as A ⟷ I ⟷ B. At 15 K/h, thermal transitions for PlyCA are observed at 41.4°C and 44.6°C, with respective Δ*H*_vH_ values of 845 kJ/mol and 674 kJ/mol. At 60 K/h, PlyCA thermal transitions are observed at 42.0°C and 48.6°C, with Δ*H*_vH_ values of 302 kJ/mol and 545 kJ/mol, respectively. The sum of the van’t Hoff enthalpies at each scan rate equals the calorimetric enthalpy, thereby indicating the thermodynamic domains of PlyCA unfold independently at these temperatures. Changes in heat capacity were not observed after cooling and rescanning the heated samples, suggesting the unfolding of PlyCA is irreversible following thermal denaturation. As shown in Fig. 4A, the exothermic event corresponding to the irreversible aggregation of PlyCA occurs immediately after the I ⟷ B thermal transition.

**Figure 4.**
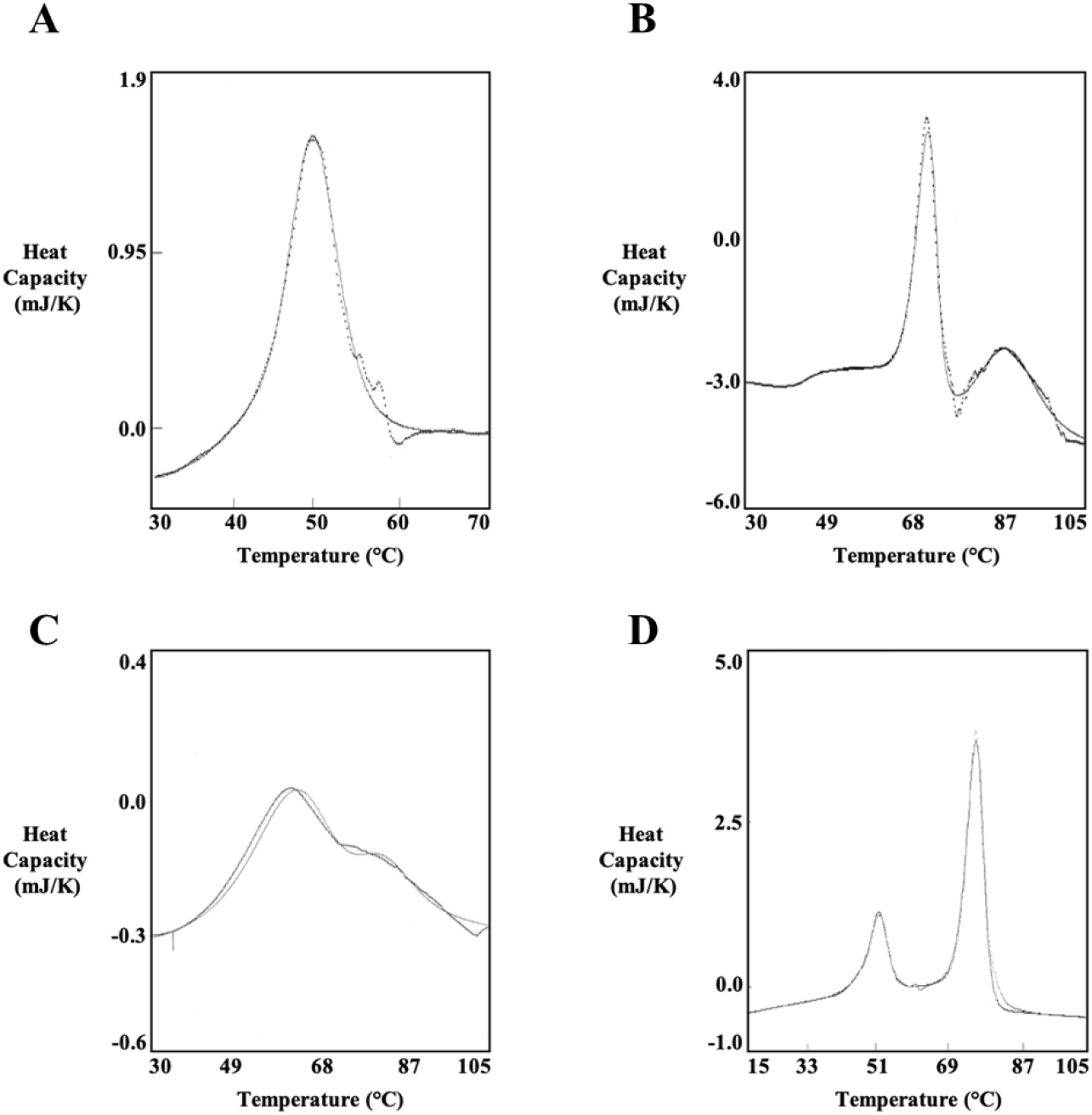
Thermodynamic analysis of PlyCA, PlyCB CBD and the PlyC holoenzyme using DSC. The thermodynamic properties of (A) PlyCA, (B) PlyCB CBD, (C) PlyCB CBD (re- scanned) and (D) PlyC were experimentally determined using DSC. Each sample was heated in the calorimeter from 15 to 105°C and subsequently cooled from 105 to 15°C using scan rates of 15 or 60 K/h. The representative thermograms depicted were obtained from analyzing the protein samples in either PBS (PlyCA and PlyC) or phosphate buffer (PlyCB CBD).

**Table 1.** Thermodynamic characterization of PlyCA in PBS by means of DSC.

| Conc.<br>(mg/ml) | Scan Rate<br>(K/h) | Model | $T_G$ (°C) | $\Delta H_{vH}$<br>(kJ/mol) | $\Delta H_{cal}$<br>(kJ/mol) | Ratio<sup>a</sup> |
| --- | --- | --- | --- | --- | --- | --- |
| 1.0 - 2.0 | 15 | A ↔ I | 41.4±1.7 | 845±154 | 1633±163 | 1.08±0.25 |
| | | I ↔ B | 44.6±0.5 | 674±194 | | |
| 1.0 – 2.0 | 60 | A ↔ I | 42.0±2.3 | 302±76 | 922±205 | 1.09±0.26 |
| | | I ↔ B | 48.6±0.2 | 545±81 | | |

<sup>a</sup>Ratio equates to [ $\Delta H_{cal}$ / ( $\Delta H_{vH}$ (low temperature) + $\Delta H_{vH}$ (high temperature))]

Next, the C-terminal CHAP domain of PlyCA was thermodynamically investigated at 1 to 2 mg/ml in PBS using scan rates of 15 and 60 K/h (Table 2). Similar to PlyCA, the resulting thermograms of the CHAP domain were each fit with A ⟷ I and I ⟷ B two-state transitions. The three-state thermal transition model of CHAP can be annotated as A ⟷ I ⟷ B. At 15 K/h, the CHAP domain displays thermal transitions at 38.5 and 44.3°C, with respective Δ*H*_vH_ values of 464 and 583 kJ/mol. When using a scan rate of 60 K/h, thermal transitions were observed at 39.1 and 46.1°C, with corresponding Δ*H*_vH_ values of 430 and 554 kJ/mol. The ratios of the calorimetric enthalpy to the van’t Hoff enthalpies were close to 0.5, indicating the CHAP domain unfolds as a single cooperative unit. Repeating DSC scans with previously heated protein samples confirmed the unfolding of the CHAP domain is an irreversible thermodynamic process.

**Table 2.**
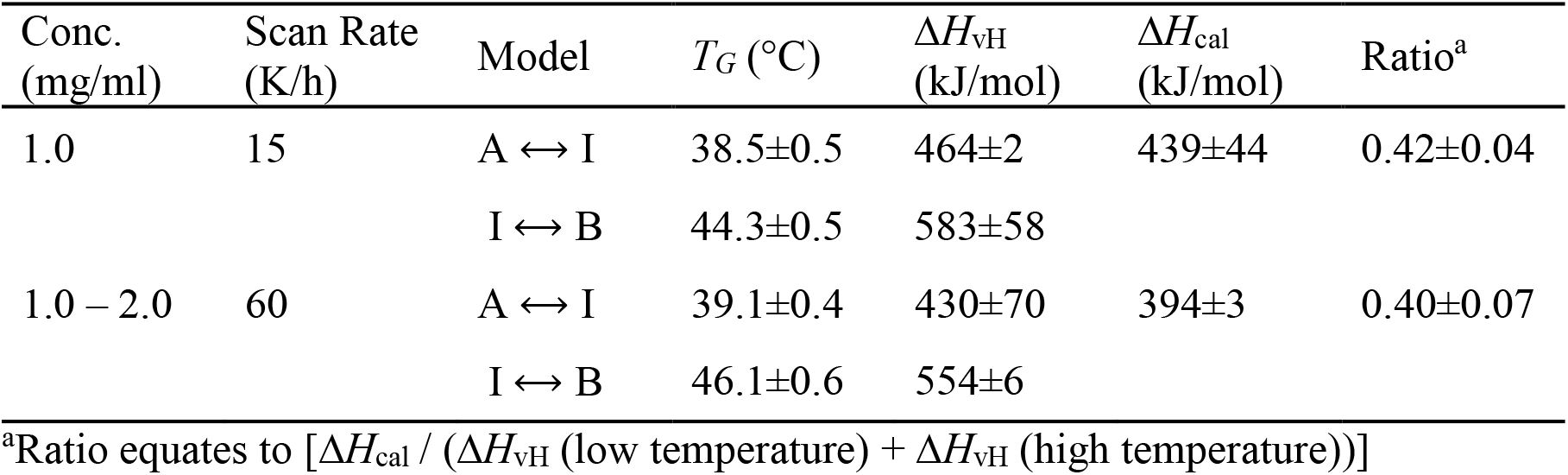
Thermodynamic characterization of the CHAP domain in PBS by means of DSC.

| Conc.<br>(mg/ml) | Scan Rate<br>(K/h) | Model | $T_G$ (°C) | $\Delta H_{vH}$<br>(kJ/mol) | $\Delta H_{cal}$<br>(kJ/mol) | Ratio<sup>a</sup> |
| --- | --- | --- | --- | --- | --- | --- |
| 1.0 | 15 | A ↔ I | 38.5±0.5 | 464±2 | 439±44 | 0.42±0.04 |
| | | I ↔ B | 44.3±0.5 | 583±58 | | |
| 1.0 – 2.0 | 60 | A ↔ I | 39.1±0.4 | 430±70 | 394±3 | 0.40±0.07 |
| | | I ↔ B | 46.1±0.6 | 554±6 | | |

<sup>a</sup>Ratio equates to [ $\Delta H_{cal} / (\Delta H_{vH} \text{ (low temperature)} + \Delta H_{vH} \text{ (high temperature)})$ ]

The thermodynamic properties of the octameric PlyCB CBD were elucidated at either 1 mg/ml in PBS (Table 3) or 1 to 2 mg/ml in phosphate buffer, pH 7.4 (Table 4). An example of a thermogram specific to the PlyCB CBD is shown in Fig. 4B. Results from the DSC scans indicate the PlyCB CBD fulfills a three-state thermal transition model summarized as 2A ⟷ 2I ⟷ 2B. The lower temperature thermal transition was fit with a two-state 2A ⟷ 2I model. This transition occurs either at 73.7°C with a Δ*H*_vH_ value of 1265 kJ/mol in PBS, or at 70.4°C with a Δ*H*_vH_ value of 1160 kJ/mol in phosphate buffer. The ratio of the transition enthalpy to the van’t Hoff enthalpy for the 2A ⟷ 2I transition was nearly 2, indicating the PlyCB octamer dissociates into an intermediate state. This intermediate state is followed by denaturation of the PlyCB subunit at the higher temperature transition. This thermal transition, which was fit with a two-state 2I ⟷ 2B model, was observed at 91.8°C with a Δ*H*_vH_ value of 368 kJ/mol in PBS, and at 83.5°C with a Δ*H*_vH_ value of 373 kJ/mol in phosphate buffer. Thus, for the 2A ⟷ 2I ⟷ 2B thermal transition model of the isolated PlyCB CBD, the first thermal transition corresponds to the dissociation of the PlyCB octamer into a monomeric intermediate state, while the second thermal transition represents the unfolding of the individual PlyCB monomers. Results from rescanning previously heated PlyCB samples indicate both thermal transitions are reversible. An example of a typical thermogram obtained from these experiments is depicted in Fig. 4C. However, the 2A ⟷ 2I transition appears at a lower temperature due to the significant reduction of folded PlyCB subunits upon rescanning, which in turn decreases the amount of protein that re-associates in the native octameric state. Additionally, the van’t Hoff enthalpy specific to the 2A ⟷ 2I transition of PlyCB was lower when rescanning the protein samples, perhaps due to the steep dependence of the van’t Hoff enthalpies on temperature.

**Table 3.** Thermodynamic characterization of the PlyCB CBD in PBS by means of DSC.

| | Conc.<br>(mg/ml) | Scan Rate<br>(K/h) | Model | $T_G$ (°C) | $\Delta H_{vH}$<br>(kJ/mol) | $\Delta H_{cal}$<br>(kJ/mol) | Ratio<sup>a</sup> |
| --- | --- | --- | --- | --- | --- | --- | --- |
| | 1.0 | 60 | 2A ↔ 2I | 73.7±0.5 | 1265±53 | 2371±237 | 1.87±0.21 |
| | | | 2I ↔ 2B | 91.8±0.1 | 368±37 | | |
| Rescan to<br>95°C | 1.0 | 60 | 2A ↔ 2I | 49.4±0.1 | 275±4 | 93±10 | |
| | | | 2I ↔ 2B | 67.0±0.1 | 230±23 | 275±28 | |
| Rescan to<br>105°C | 1.0 | 60 | 2A ↔ 2I | 68.2±0.1 | 263±26 | 148±15 | |
| | | | 2I ↔ 2B | 91.5±0.1 | 650±65 | 57±6 | |

<sup>a</sup>Ratio equates to [ $\Delta H_{cal}$ / ( $\Delta H_{vH}$ (low temperature))]

**Table 4.** Thermodynamic characterization of the PlyCB CBD in phosphate buffer by means of DSC.

| | Conc.<br>(mg/ml) | Scan Rate<br>(K/h) | Model | $T_G$ (°C) | $\Delta H_{vH}$<br>(kJ/mol) | $\Delta H_{cal}$<br>(kJ/mol) | Ratio<sup>a</sup> |
| --- | --- | --- | --- | --- | --- | --- | --- |
| | 1.0 – 2.0 | 60 | 2A ↔ 2I | 70.4±1.4 | 1160±119 | 2349±235 | 2.03±0.28 |
| | | | 2I ↔ 2B | 83.5±2.7 | 373±84 | | |
| Rescan to<br>75°C | 1.0 | 60 | 2A ↔ 2B | 68.8±0.1 | 945±95 | 1870±187 | |
| Rescan to<br>95°C | 2.0 | 60 | 2A ↔ 2I | 63.0±0.1 | 427±43 | 281±28 | |
| | | | 2I ↔ 2B | 82.4±0.1 | 405±13 | | |
| Rescan to<br>105°C | 1.0 | 60 | 2A ↔ 2I | 58.1±0.1 | 241±24 | 555±56 | |
| | | | 2I ↔ 2B | 82.1±0.1 | 227±3 | | |

<sup>a</sup>Ratio equates to $[\Delta H_{cal} / (\Delta H_{vH} \text{ (low temperature)})]$

DSC scans of the PlyC holoenzyme were performed at concentrations ranging from 0.81 to 3.82 mg/ml in PBS using scan rates of 15 and 60 K/h (Table 5). An example of a DSC thermogram of PlyC is shown in Fig. 4D. The complete thermal denaturation of PlyC satisfies a four-state thermal transition model of A ⟷ I ⟷ B for the PlyCA subunit and 2A ⟷ 2B for the PlyCB CBD. At 15 K/h, the first two thermal transitions observed at 44.5 and 48.4°C, which had respective Δ*H*_vH_ values of 677 and 862 kJ/mol, were fit with A ⟷ I and I ⟷ B two-state transitions. The third thermal transition at 73.7°C, which displayed a Δ*H*_vH_ value of 1238 kJ/mol, was fit with a 2A ⟷ 2B two-state model. When using a scan rate of 60 K/h, the initial two thermal transitions are seen at 46.2 and 50.6°C, with respective Δ*H*_vH_ values of 472 and 637 kJ/mol, were fit with A ⟷ I and I ⟷ B two-state transitions. The third thermal transition at 75.0°C, which displayed a Δ*H*_vH_ value of 1082 kJ/mol, was fit with a 2A ⟷ 2B two-state model. The lower temperature thermal transitions of PlyC at both scan rates correlate to the thermal denaturation of PlyCA. The ratio of the calorimetric enthalpy to the van’t Hoff enthalpies is nearly 0.5, suggesting the GyH and CHAP domains of PlyCA are unfolding cooperatively. The higher temperature thermal transition is specific to the thermal denaturation of the PlyCB CBD. This thermal transition equates to the 2A ⟷ 2I transition displayed when scanning the PlyCB CBD alone. The 2I ⟷ 2B transition at 91.8°C observed with PlyCB alone is not observed when scanning the holoenzyme. Similar to DSC experiments using PlyCA and PlyCB CBD alone, cooling and rescanning previously heated PlyC samples revealed the respective irreversible and reversible folding characteristics of PlyCA and the PlyCB CBD. Only a small subpopulation of PlyCB subunits refolded, resulting in a decreased thermal transition temperature and van’t Hoff enthalpy when compared to the initial DSC scan.

**Table 5.**
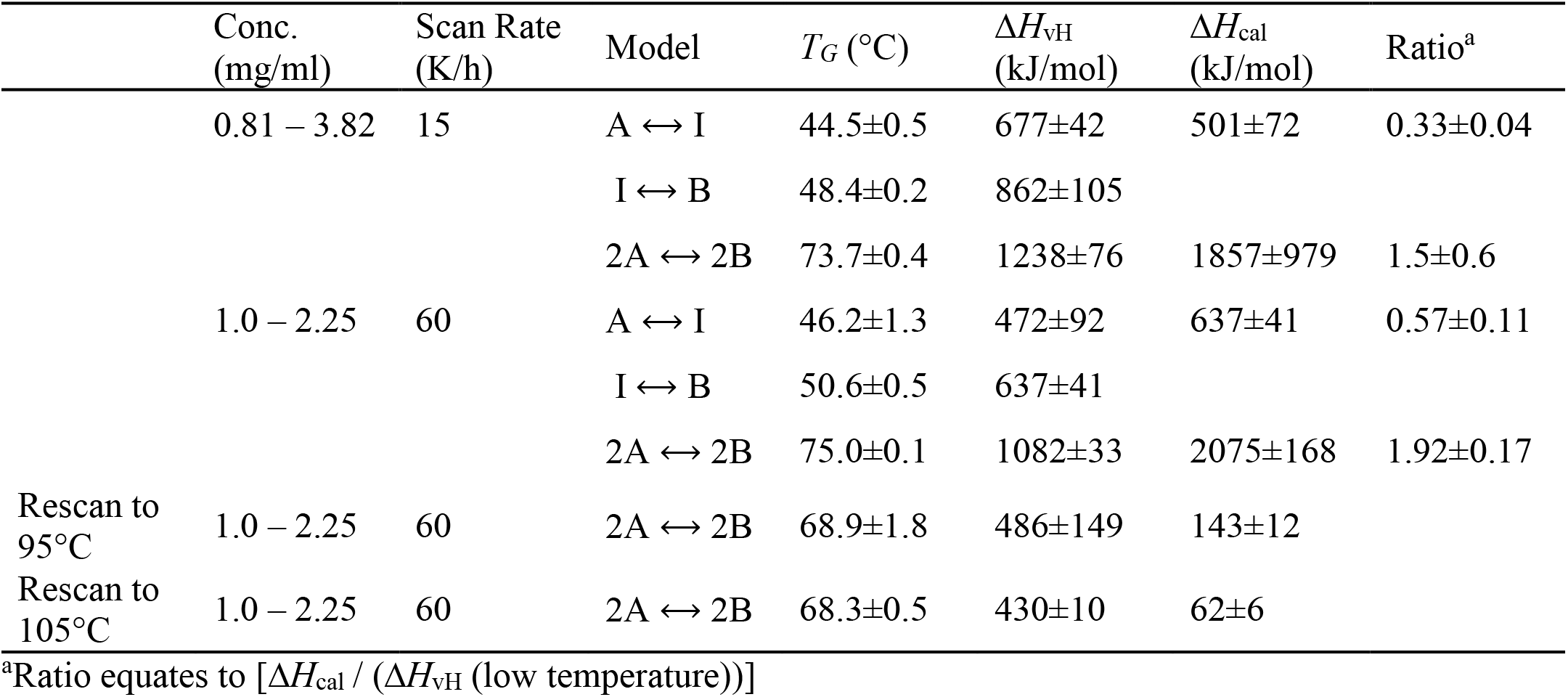
Thermodynamic characterization of PlyC in PBS by means of DSC.

| | Conc.<br>(mg/ml) | Scan Rate<br>(K/h) | Model | $T_G$ (°C) | $\Delta H_{vH}$<br>(kJ/mol) | $\Delta H_{cal}$<br>(kJ/mol) | Ratio<sup>a</sup> |
| --- | --- | --- | --- | --- | --- | --- | --- |
| | 0.81 – 3.82 | 15 | A ↔ I | 44.5±0.5 | 677±42 | 501±72 | 0.33±0.04 |
| | | | I ↔ B | 48.4±0.2 | 862±105 | | |
| | | | 2A ↔ 2B | 73.7±0.4 | 1238±76 | 1857±979 | 1.5±0.6 |
| | 1.0 – 2.25 | 60 | A ↔ I | 46.2±1.3 | 472±92 | 637±41 | 0.57±0.11 |
| | | | I ↔ B | 50.6±0.5 | 637±41 | | |
| | | | 2A ↔ 2B | 75.0±0.1 | 1082±33 | 2075±168 | 1.92±0.17 |
| Rescan to<br>95°C | 1.0 – 2.25 | 60 | 2A ↔ 2B | 68.9±1.8 | 486±149 | 143±12 | |
| Rescan to<br>105°C | 1.0 – 2.25 | 60 | 2A ↔ 2B | 68.3±0.5 | 430±10 | 62±6 | |

<sup>a</sup>Ratio equates to [ $\Delta H_{cal}$ / ( $\Delta H_{vH}$ (low temperature))]

### PlyCB octamer structural integrity and kinetic stability as a function of temperature

The DSC results for the isolated PlyCB CBD in PBS dictate the octamer reversibly dissociates into an intermediate state at 73.7°C, which is then followed by the individual subunits thermally denaturing at 91.8°C (Table 3). To further expand on these findings, the PlyCB CBD was heated in PBS to temperatures ranging from 65 to 90°C, followed by chemical cross-linking in order to qualitatively assess the structural integrity of the octamer via SDS-PAGE. At temperatures ≤ 74.4°C, monomers, dimers, trimers and progressively higher ordered species of PlyCB were observed (Fig. 5A). In contrast, at temperatures exceeding 74.4°C, there is a distinct change in the manner the PlyCB subunits interact, resulting in only monomers and dimers of PlyCB being observed. Consistent with the DSC findings, these results indicate the PlyCB octamer experiences a thermally-induced change in quaternary structure at temperatures ≥ 75°C.

**Figure 5.**
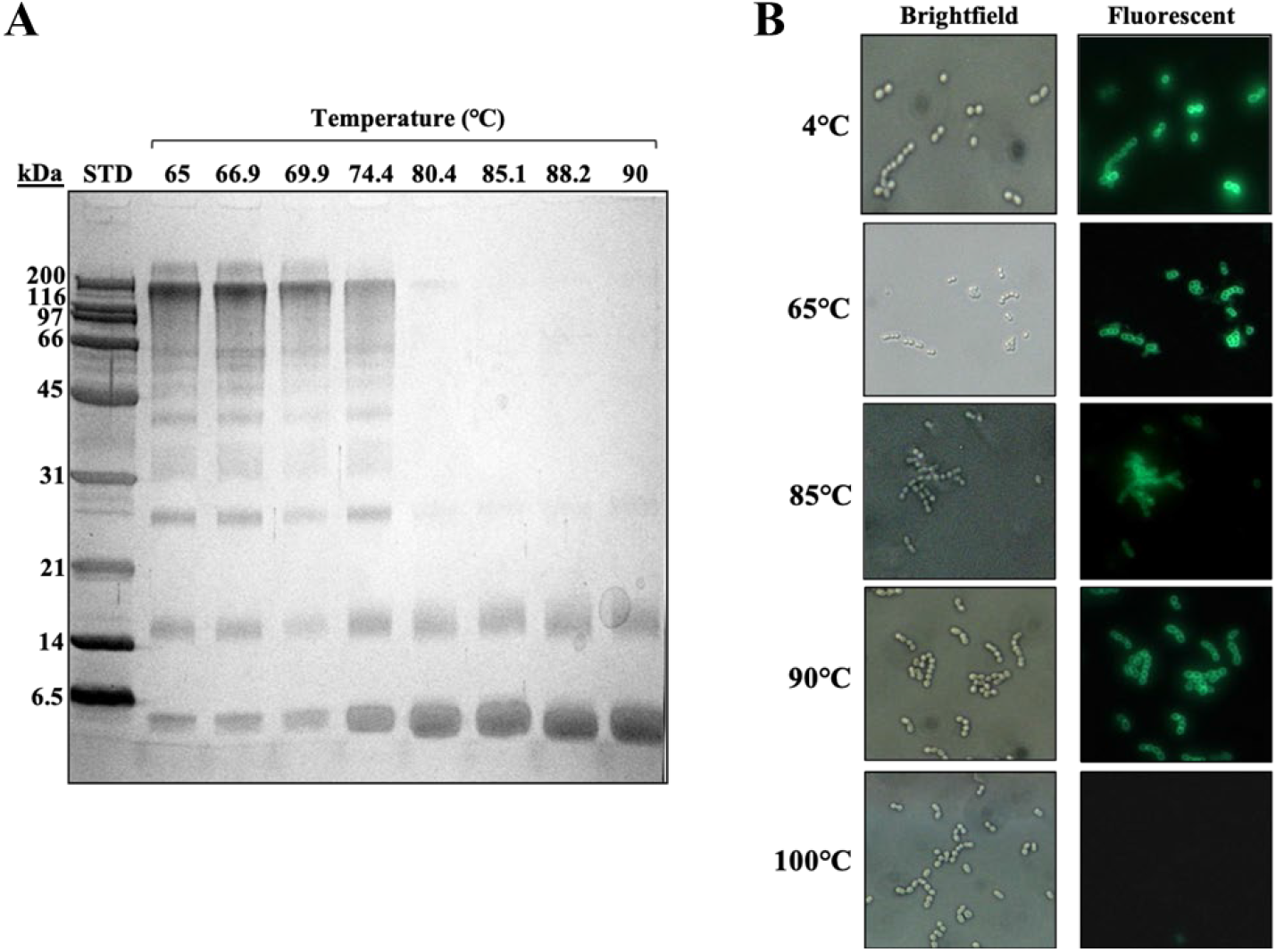
Structural and kinetic stability of the PlyCB octamer. (A) Following a 10-min incubation in PBS at temperatures ranging from 65 to 90°C, PlyCB at 1 mg/ml was chemically cross-linked and cooled on ice. The samples were analyzed using SDS-PAGE in order to evaluate the structural integrity of the PlyCB octamer. (B) AlexaFluor-labeled PlyCB CBD at 1 mg/ml in PBS was incubated at 4 to 100°C for 30 min, cooled on ice, and then added to *S. pyogenes*. The kinetic stability of the heat-treated PlyCB samples (i.e., PlyCB that retains its ability to bind *S. pyogenes* following heat treatment) was analyzed by means of fluorescence microscopy.

When incubating the PlyCB CBD at temperatures below its thermal denaturation temperature, the reversibility of its intermediate state implies the octamer should be able to re-associate upon cooling and thus retain its ability to bind target streptococci. To this end, fluorescently-labeled PlyCB CBD was incubated in PBS at temperatures ranging from 4 to 100°C, cooled and then assayed for binding to *S. pyogenes* using fluorescence microscopy (Fig. 5B). At temperatures ≤ 90°C, the PlyCB CBD is capable of either maintaining or refolding into its octameric structure in order to bind the streptococcal surface. Alternatively, incubation at 100°C inhibits PlyCB functionality due to the thermal denaturation of its subunits. This is functional evidence the intermediate state of PlyCB is reversible.

### Refolding of the PlyC holoenzyme

With the dissociation temperature and the reversibility of the intermediate state of PlyCB being experimentally validated, the folding dynamics of the PlyC holoenzyme were investigated next as a function of the PlyCB quaternary structure. A series of functional assays were performed to determine if PlyC can be reconstituted by mixing PlyCA with either PlyCB in its octameric state or with individual PlyCB subunits concurrently oligomerizing. For the initial set of control experiments, equal molar concentrations of PlyC, PlyCA or PlyCB CBD were added to *S. pyogenes* and the bacteriolytic activity was subsequently assessed using turbidity reduction assays (Fig. 6). Data was normalized to the lytic activity displayed by native PlyC. As expected, in the absence of the PlyCA catalytic subunit, the PlyCB CBD displays no lytic activity towards *S. pyogenes*. Furthermore, without the PlyCB CBD to direct the catalytic subunit to its peptidoglycan substrate, PlyCA alone exhibited < 1% in relative activity compared to the holoenzyme. Next, using a molar ratio of approximately 1:1, PlyCA was mixed with the PlyCB CBD at room temperature. The bacteriolytic activity displayed by the resulting mixture was equal to that of PlyCA alone, suggesting the structure of the PlyC holoenzyme was not regenerated. With this understanding, a near equal molar concentration of ice-cold PlyCA was rapidly mixed with PlyCB that had been heated to 75°C, and the mixture was then rapidly cooled. In this scenario, the PlyCB octamer dissociates into an intermediate state when heated and then promptly re-associates when PlyCA is added and the mixture is cooled. When added to *S. pyogenes*, the reconstituted enzyme retained 35% of the relative bacteriolytic activity of native PlyC. Results from gel filtration indicated the aforementioned reconstituted enzyme had a molecular mass of ∼120 kDa (data not shown), which equates to the mass of the PlyC holoenzyme. This is both functional and biochemical evidence that PlyCA is interacting with the PlyCB octamer to regenerate the PlyC holoenzyme structure.

**Figure 6.**
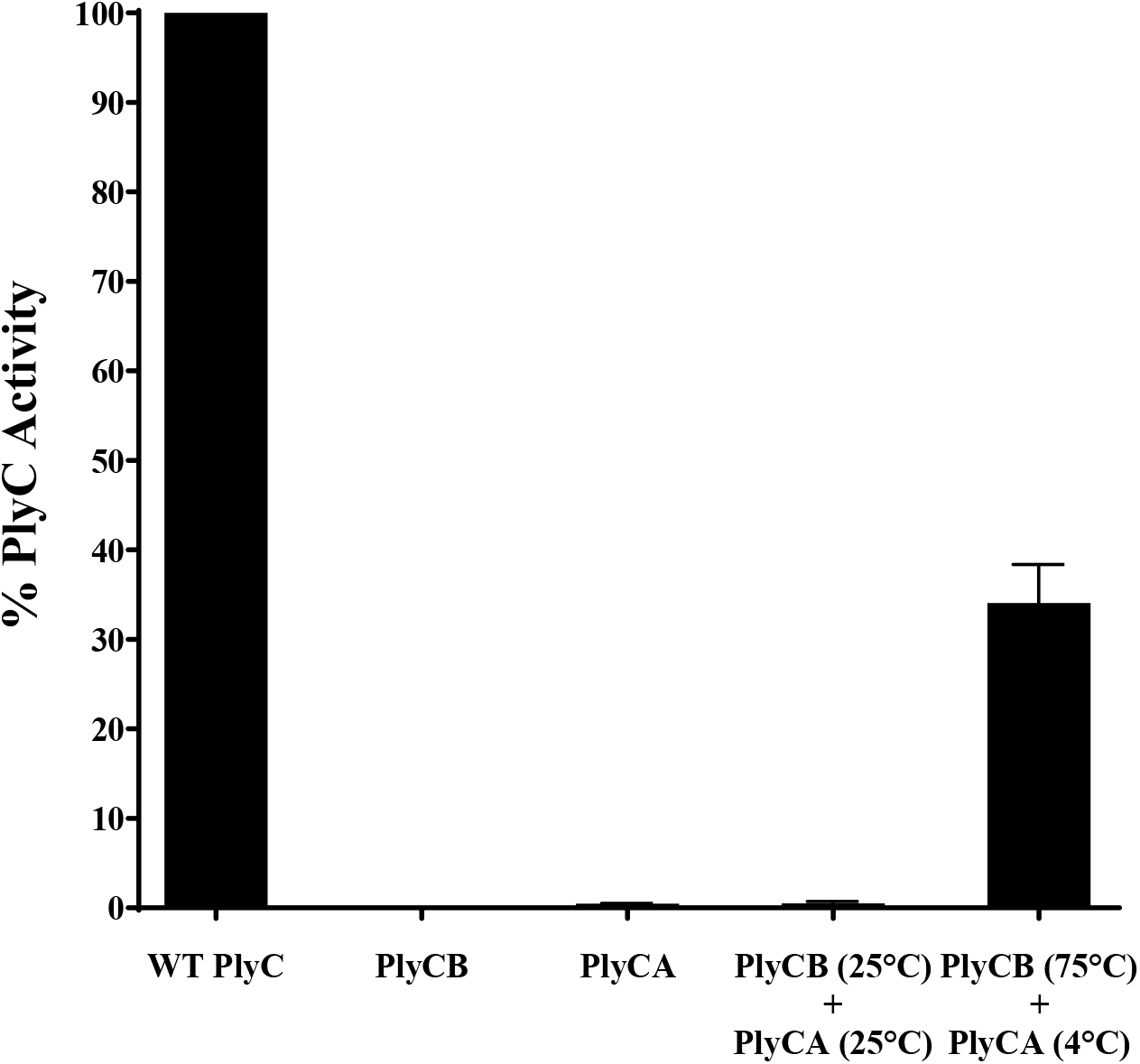
Thermal reconstitution of the PlyC holoenzyme. To elucidate the folding dynamics of PlyC, an attempt was made to reassemble the holoenzyme structure using purified PlyCA and PlyCB subunits. In one scenario, PlyCA and the PlyCB CBD were incubated at room temperature. Alternatively, ice-cold PlyCA was mixed with the PlyCB CBD in PBS heated to 75°C and then immediately placed on ice. A 1:1 molar ratio of PlyCA:PlyCB CBD was used in both instances. Used as an indirect measure of PlyC holoenzyme reconstitution, the bacteriolytic activity of each sample at equal molar concentrations was assayed against *S. pyogenes* using a turbidity reduction assay. Activity was normalized to that of PlyC. Isolated PlyCA (< 1% activity of PlyC) and PlyCB CBD (no activity) were used as controls.

## Discussion

There is currently a deficiency in studies directed towards obtaining a thorough understanding of the thermal characteristics of bacteriophage endolysins. Results from such analyses would allow for a more complete and accurate assessment of the therapeutic potential of the biologic, as well as provide insightful information for downstream development. Previous studies evaluating the thermal properties of PlyC revealed the enzyme has modest kinetic stability (19–21). This finding seemed to be attributed to the thermosusceptiblility of PlyCA. To further confirm this observation, an extensive thermal characterization of PlyC was performed. When isolating each component, DSC analysis in PBS showed that the thermodynamic stability of PlyCA (*T_G_* ∼42°C; Fig. 4A and Table 1) is about 32°C lower than that of the PlyCB CBD (*T_G_* ∼74°C; Table 3). Using DSC to analyze the thermodynamics of the PlyC holoenzyme revealed the PlyCA subunit unfolds initially, displaying two cooperative thermal transitions between approximately 45°C and 50°C (Fig. 4D and Table 5). As confirmed by both DSC and functional assays, PlyCA unfolding is irreversible, which consequently results in kinetic inactivation of the enzyme at 46°C in PBS (Fig. 2, bottom). Following PlyCA unfolding, the PlyCB CBD thermally denatures at ∼74°C (Fig. 4D and Table 5).

There are notable differences in the thermal transition properties of both PlyCA and the PlyCB CBD when each component is analyzed either alone or complexed as a holoenzyme. For example, the thermodynamic stability of PlyCA is enhanced by about 4°C when complexed with the PlyCB octamer (compare Tables 1 and 5). Moreover, the ratios of the calorimetric enthalpy to the sum of the van’t Hoff enthalpies specific to the two PlyCA thermal transitions are close to 0.5 and 1 in the PlyC and PlyCA only samples, respectively. These ratios indicate the GyH and CHAP domains of PlyCA can unfold independently when the catalytic subunit is uncomplexed, or as a single cooperative unit when PlyCA is anchored to the PlyCB CBD in the PlyC holoenzyme. These observations signify the GyH and CHAP domains interact strongly when PlyCA is bound to the PlyCB CBD, which is consistent with the synergistic activities noted between these two domains in the holoenzyme (11). Like PlyCA, the thermal transition properties of the PlyCB CBD vary depending on whether the octameric binding domain is analyzed alone or complexed with the catalytic subunit. DSC scans of the PlyCB CBD alone in PBS shows two uncooperative thermal transitions at 74°C (octamer dissociation) and 92°C (PlyCB monomer unfolding) (Table 3). Thermal cross-linking experiments supported these results, with the PlyCB octamer appearing to dissociate at temperatures ≥ 75°C (Fig. 5A). Additionally, heating/cooling experiments coupled with fluorescence microscopy confirmed that PlyCB can maintain or refold into its octameric structure to bind *S. pyogenes* until heated to 100°C (Fig. 5B). While the lower thermal transition of PlyCB is maintained during DSC scans of PlyC, the higher 2I ⟷ 2B transition at 92°C is absent (Table 5). This result suggests PlyCB unfolding in the context of the holoenzyme does not proceed through an intermediate state. Instead, it appears the PlyCB monomers simultaneously unfold as the octamer dissociates.

When analyzing the calorimetric data for the PlyCB CBD in PBS (Table 3) and phosphate buffer (Table 4), it is evident that buffer composition has a significant effect on stability. Compared to phosphate buffer, the *T*_G_ for the 2A ⟷ 2I and 2I ⟷ 2B thermal transitions of the PlyCB CBD was increased by a respective 3.3 and 8.3°C in PBS. The thermodynamic stability of both the octamer and the individual PlyCB monomers appear to be enhanced by the presence of salt. As seen with other proteins (e.g., (22–24)), optimizing surface charge-charge networks by displacing water with electrostatic interactions between salts and proteins can improve stability (25). For the PlyCB CBD, a direct relationship between salt concentration and protein stability can be assayed by performing future DSC experiments in phosphate buffer supplemented with varying concentrations of NaCl.

The crystal structure of the uncomplexed PlyCB octamer shows disordered electron density for the eight N-terminal residues of each PlyCB monomer (11), presumably due to flexibility of this region. However, in the crystal structure of the PlyC holoenzyme, these same N-terminal amino acids form at least a four-stranded parallel β-sheet that creates a platform upon which the PlyCA central docking domain interacts via an antiparallel bundle of three α-helices. These data suggest PlyCA and PlyCB fold together at the time of translation. Indeed, co-expression of PlyCA and PlyCB on separate plasmids allows for the reconstitution of a functional PlyC holoenzyme structure (26), yet mixing pre-formed PlyCA and PlyCB CBD at room temperature does not yield the active holoenzyme (Fig. 6). We therefore sought to elucidate whether the pre-folded state of the PlyCB CBD could be simulated based on its unique thermodynamic properties in order to study the folding dynamics of PlyC. In its uncomplexed form, the thermally-induced reversible folding of the PlyCB octamer allowed for the *in vitro* reconstitution of the PlyC holoenzyme. This was accomplished by adding ice-cold PlyCA to PlyCB in a heated intermediate state, and subsequently cooling the mixture on ice (Fig. 6). These findings indicate that the holoenzyme forms by PlyCA complexing with individual PlyCB subunits as they are simultaneously self-associating into the octameric ring structure.

There is preliminary evidence to support that the CHAP domain of PlyCA is the most thermosusceptible structural component of PlyC. The isolated CHAP domain unfolds at 39°C (Table 2), which is a 3°C decrease from that of the isolated full-length PlyCA subunit (Table 1). Additional PlyC constructs will need to be thermodynamically investigated to definitively prove whether the CHAP domain is indeed the least thermostable component of the holoenzyme. Such constructs would include PlyCΔCHAP (i.e., PlyC holoenzyme without the CHAP domain of PlyCA), the isolated GyH domain of PlyCA, PlyCΔGyH (i.e., PlyC holoenzyme without the GyH domain of PlyCA) and PlyCΔΔ (i.e., the PlyCB CBD complexed with the helical docking domain of PlyCA only).

The overall findings from this study indicate the PlyCB CBD is highly thermostable, while the PlyCA subunit is relatively thermolabile. This general understanding will aid in formulating future engineering strategies for enhancing the thermal stability of PlyC. For example, PlyCA can be subjected to an optimized directed evolution-based methodology specific to increasing the thermal stability of bacteriolytic enzymes; this method was validated using PlyC (21). Alternatively, the PlyC crystal structure allows for structure-guided engineering approaches to be employed. Structural characteristics conserved in thermophilic proteins can be incorporated into PlyCA using rationale-based site-directed mutagenesis, such as substituting uncharged polar amino acids with charged residues to increase surface charge-charge interactions, shortening loop and turn structures, reducing the occurrence of thermolabile residues, and decreasing the size and amount of hydrophobic core cavities (27). As shown in previous studies, intra- and inter-domain disulfide bonds can be introduced to decrease the entropy associated with the folded or unfolded state of the protein, thereby increasing stability (28, 29). Computational methods utilizing *in silico* protein folding algorithms can also be used to estimate free energy values of folding, with a goal of predicting point mutations that are thermally advantageous to PlyCA. This strategy has already been proven useful for increasing the thermal stability of endolysins, specifically PlyC. Applying the Rosetta and FoldX protein folding algorithms to the CHAP domain of PlyCA allowed for the identification of a single point mutation that yielded a 16-fold increase in kinetic stability when compared to wild-type (20). Domain swapping is another engineering approach that can be used to increase the thermal stability of PlyC. For example, due to its modular structure, the CHAP domain of PlyCA can be substituted with a catalytic domain originating from an intrinsically thermostable peptidoglycan hydrolase. It is worth noting, however, that this type of engineering approach may not be optimal for a complex endolysin like PlyC, as the synergism shared between the GyH and CHAP domains may be disrupted and generate significant activity defects.

## Conclusions

The thermolabile nature of the bacteriolytic mechanism of PlyC has been understood since the early 1970s (19). An in-depth thermal characterization has been lacking to identify the structural component(s) of the holoenzyme that is/are responsible for the modest stability of the endolysin. Results from our studies revealed the PlyCB CBD of PlyC is highly thermostable, whereas the PlyCA catalytic subunit is reasonably thermosusceptible. We speculate that the poor intrinsic stability of PlyCA could be due to its C-terminal CHAP domain, which was shown to be the most thermodynamically unstable PlyC-related construct analyzed. However, this result is preliminary, as additional PlyC constructs focusing on the GyH and helical docking domains of PlyCA need to be designed and studied using DSC.

There were significant differences in the thermal transition properties of the PlyC holoenzyme compared to its isolated catalytic subunit and CBD. For example, the GyH and CHAP domains of PlyCA appear to unfold independently when the catalytic subunit is isolated. When PlyCA is complexed to the PlyCB octamer, however, the GyH and CHAP domains interact strongly and unfold cooperatively. The thermodynamic stability of the PlyCA subunit was also enhanced when bound to the PlyCB CBD. In the context of the PlyC holoenzyme, the PlyCB CBD undergoes a single thermal transition corresponding to the simultaneous dissociation of the octamer, as well as each subunit denaturing. When isolated, the PlyCB octamer dissociates into a reversible intermediate state, which is then followed by a higher temperature thermal transition relating to each subunit denaturing. Adding ice-cold PlyCA to the intermediate state of PlyCB, followed by rapid cooling, resulted in the reconstitution of an enzymatically-active holoenzyme structure. This observation could aid in understanding the *in vivo* folding dynamics of this complex, multimeric endolysin.

## Conflict of interest

DCN is co-inventor of patents related to the PlyC endolysin.

## Acknowledgements

We would like to thank the National Institute of Standards and Technology for sharing their VP-DSC Microcalorimeter. This manuscript is dedicated to our late co-author, Frederick P. Schwarz.

## Author contributions

**JTH:** Conceptualization, Data curation, Formal analysis, Writing – original draft. **RDH:** Conceptualization, Data curation, Formal analysis, Validation, Writing – review and editing. **FPS:** Conceptualization, Data curation, Formal analysis, Writing – original draft. **DCN:** Project administration, Funding acquisition, Formal analysis, Writing – review and editing.

## Notes

### Summary of Updates

New data has been added. Specifically, a new Figure 1 has been added on the kinetic stability of the enzymes. Additional edits to the text have been made throughout the manuscript to provide more clarity.

